# Sex-based *de novo* transcriptome assemblies of the parasitoid wasp *Encarsia suzannae*, a host of the manipulative heritable symbiont *Cardinium hertigii*

**DOI:** 10.1101/2022.08.05.502955

**Authors:** Dylan L Schultz, Evelyne Selberherr, Corinne M Stouthamer, Matthew R Doremus, Suzanne E Kelly, Martha S Hunter, Stephan Schmitz-Esser

## Abstract

Minute parasitoid wasps in the genus *Encarsia* are commonly used as biological pest control agents of whiteflies and armored scale insects in greenhouses or in the field. They are also a key host of the bacterial endosymbiont *Cardinium hertigii* which can cause a suite of reproductive manipulation phenotypes, including parthenogenesis, feminization, and cytoplasmic incompatibility; the last being most thoroughly studied in *Encarsia suzannae*. Despite their biological and economic importance, there are currently no published *Encarsia* genomes and only one public transcriptome. In this study, we applied a mapping-and-removal approach to eliminate known contaminants from previously-obtained Illumina sequencing data. We generated *de novo* transcriptome assemblies for both female and male *E. suzannae* which contain 45,986 and 54,762 final coding sequences, respectively. Benchmarking Single-Copy Orthologs (BUSCO) results indicate both assemblies are highly complete. Preliminary analyses revealed the presence of homologs of sex-determination genes characterized in other insects and putative venom proteins. These transcriptomes will be valuable tools to better understand the biology of *Encarsia* wasps and their evolutionary relatives. Furthermore, the separate male and female assemblies will be particularly useful references for studies involving insects of only one sex.

## Background

*Encarsia suzannae* are minute parasitoid wasps within the order Hymenoptera and are of interest due to their unusual behavior and biology, their use in biological control of the important whitefly pest *Bemisia tabaci*, their relatedness to the widespread greenhouse biological control agent *Encarsia formosa*, and because they harbor a bacterial endosymbiont capable of host reproductive manipulation, *Cardinium hertigii. Cardinium*, in the bacterial phylum *Bacteroidota*, shows independent evolution of reproductive manipulation from the well-known alphaproteobacterial *Wolbachia* [1]. Like other Hymenoptera, *E. suzannae* are haplodiploid and reproduce via arrhenotoky (arrhenotokous parthenogenesis) in which haploid males are produced via unfertilized eggs and females are derived from fertilized diploid eggs [2]. Unusually, most *Encarsia* species, including *E. suzannae*, are also autoparasitoids, with females developing in and consuming the nymphs of the sweet potato whitefly, *B. tabaci*, while male wasps develop as hyperparasitoids, consuming the pupae of conspecific females or other aphelinid parasitoids of whiteflies. After consuming their host, both male and female *Encarsia* pupate in the whitefly cuticle and emerge as adults [3]. Many *Encarsia* species are effective parasites of whitefly species, which are widespread pests causing up to billions of dollars in crop losses yearly as they can directly damage plants by feeding and are able to transmit more than 200 different plant viruses to a multitude of plant species [4, 5]. As a result, *Encarsia* species have been widely used as pest control agents to limit whitefly populations in field or greenhouse settings [6-8]. Their unusual autoparasitic biology [9], sex allocation behavior, and host selection have also been the focus of study in these intriguing wasps [10].

Like many insects, *Encarsia* may be infected with maternally-transmitted intracellular bacterial endosymbionts, such as *Wolbachia* and *Cardinium*, which influence their transmission by manipulating host reproduction [11] or oviposition behavior [12] to favor infected females. These manipulations include the induction of asexual reproduction via thelytokous parthenogenesis [13, 14], as well as a type of male reproductive sabotage called cytoplasmic incompatibility (CI) [15]. CI causes the offspring of infected males and uninfected females to die early in development, yet females infected with the same symbiont can successfully mate with infected or uninfected males. This sabotage proceeds via a two-step mechanism in which the symbiont alters male sperm with a fatal modification, then rescues infected offspring from this fatal modification when present in the fertilized egg. Together, the modification and rescue steps of CI grant infected females with a relative fitness advantage over uninfected females, which drives the symbiont to high frequencies in host populations [11]. The role of endosymbionts in arthropod biology, evolution, and speciation have been a subject of intense study [16-18]. Much of this research has focused on symbiont-induced CI given its potential role in insect speciation [19-21], its application in arthropod pest population control [22, 23], and its ability to drive desirable genetic traits through populations (e.g. resistance to arthropod-borne diseases) [24].

The *c*Eper1 strain of *Cardinium hertigii* is the causal agent of CI in *E. suzannae* [15]. This symbiosis between *c*Eper1 and *E. suzannae* is the best-studied instance of *Cardinium*-induced CI, and this strain of *Cardinium* has been well-characterized by genomic and transcriptomic data [1, 3]. However, sequence information on the host, *E. suzannae*, is extremely sparse: it currently lacks a sequenced genome and a transcriptomic profile, hampering the molecular identification of host-symbiont interactions. Here, we have generated separate *de novo* assembled transcriptomes for male and female *E. suzannae* using previously obtained RNA-seq data generated to characterize the *Cardinium hertigii* transcriptome [3]. To our knowledge, there is only one other publicly available *Encarsia* transcriptome: that of the widely used greenhouse whitefly biocontrol agent *Encarsia formosa*, which has been published as part of a phylogenetic characterization of Chalcidoidea parasitoid wasps [25, 26]. Based on differences in morphology and lifestyle between *E. suzannae* and *E. formosa*, as well as their phylogenetic relationship, the two species are distantly related within the diverse *Encarsia* genus [27-30]. This dataset will be a valuable asset in an ecologically important lineage of chalcidoid wasps (Aphelinidae) that is sorely lacking sequencing data, as well as provide the first molecular characterization of the host in the model *Cardinium* CI system.

## Methods

### Sample information and sequencing

Transcriptome data was obtained as described by Mann et. al [3] which considered only *Cardinium* reads. Here, we focused on the host (non-*Cardinium*) reads from that dataset. In brief, the initial *E. suzannae* culture was obtained in 2006 in Weslaco, TX from whitefly (*B. tabaci)* hosts. Male and female wasps were reared separately in a laboratory culture as described previously [3]. For females, mated *E. suzannae* were introduced to cages bearing whitefly nymphs on cowpea (*Vigna unguiculata)* plants. For males, unmated *E. suzannae* were provided with *Eretmocerus*. sp. nr. *emiratus* larvae or pupae developing within whitefly nymphs. Total RNA from 6 groups of 350-500 male or female 1-to 3-day old *E. suzannae* wasps was extracted using the Trizol reagent (Invitrogen) followed by digestion of genomic DNA with the Turbo DNA-free kit (Ambion). The quality of extracted RNA was assessed with an Agilent 2100 bioanalyzer (Agilent Technologies) and three libraries for each sex were generated with the NEBNext Ultra RNA library prep kit combined with rRNA depletion via the Ribo-Zero Magnetic Gold kit (Epicentre Biotechnologies). Samples were sequenced on an Illumina HiSeq2500 platform at the Vienna BioCenter Core Facilities (VBCF) NGS unit [31], producing a range of 127 to 162 million 50bp paired-end reads per sample [3].

### Read preparation and assembly

Raw read files were processed with BBDuk from the BBTools suite of software (v37.36) [32] to remove Illumina adapter sequences, trim and/or filter out whole reads with a quality score less than 15, and remove reads shorter than 36bp after trimming via the following options: “ref=adapters.fa ktrim=r ordered k=23 hdist=1 mink=11 tpe tbo maq=15 qtrim=rl trimq=15 minlen=36”. We utilized FastQC (v0.11.9) to visualize the sequence quality of each sample before and after trimming and to confirm the successful removal of adapter sequences [33]. Due to the complex biology of this species and its host insects, sequence contamination from a variety of organisms throughout the rearing system is inevitable, including *Cardinium c*Eper1, the different insect hosts of male and female *E. suzannae*, and the endosymbionts of those insect hosts. Thus, we employed a mapping-and-removal approach to enrich for *E. suzannae* reads prior to assembly and limit the generation of contaminating transcripts. In this approach, BBMap (from BBTools) was used to initially map quality-trimmed reads to the genomes of *Cardinium hertigii c*Eper1 and the endosymbionts of *Bemisia tabaci* MEAM1 (with which *E. suzannae* females and males have direct or indirect contact): *Hamiltonella defensa, Portiera aleyrodidarum*, and *Rickettsia* sp. MEAM1 [34, 35]. It was also determined that the *E*. sp. nr. *emiratus* hosts of *E. suzannae* males contain *Wolbachia* [36]; thus, the *Wolbachia* wPip genome was added and mapped to the male samples. Reads that did not map to any of these bacterial genomes with greater than 94% identity (to allow for a difference of 3 nucleotides between sequenced transcripts and reference endosymbiont genomes) were retained. These reads were then subsequently mapped to the *B. tabaci* MEAM1 genome with a more stringent 97% identity threshold using BBMap to avoid mapping *E. suzannae* reads from genes highly conserved in both *Encarsia* and *Bemisia* (see Table 1 for mapping and removal details). Again, only unmapped reads were retained for assembly, as these final reads are expected to be mainly attributed to *E. suzannae*.

**Table 1:** Pre-assembly contaminant read mapping and removal of *Encarsia suzannae* transcriptome sequencing data.

| Organism | Reason for removal | Proportion of trimmed reads mapped | GenBank accession no. |
| --- | --- | --- | --- |
| *Cardinium hertigii* cEper1 | CI-causing secondary *E. suzannae* endosymbiont | <u>Female</u>: 1.183 %<br><u>Male</u>: 0.991 % | GCA_000304455.1 |
| *Portiera aleyrodidarum* MEAM1 | Primary endosymbiont of *B. tabaci* | <u>Female</u>: 0.043 %<br><u>Male</u>: 0.035 % | GCA_002285875.1 |
| *Rickettsia* sp. MEAM1 | Secondary endosymbiont of *B. tabaci* | <u>Female</u>: 0.058 %<br><u>Male</u>: 0.065 % | GCA_002285905.1 |
| *Hamiltonella defensa* MEAM1 | Secondary endosymbiont of *B. tabaci* | <u>Female</u>: 0.040 %<br><u>Male</u>: 0.037 % | GCA_002285855.1 |
| *Bemisia tabaci* MEAM1 | Parasitized by female *E. suzannae* offspring and *E. sp. nr. emiratus* | <u>Female</u>: 5.343 %<br><u>Male</u>: 5.289 % | GCA_001854935.1 |
| *Wolbachia pipientis* wPip (male only) | Secondary endosymbiont of *E. sp. nr. emiratus* which is parasitized by male *E. suzannae* offspring | <u>Female</u>: N/A<br><u>Male</u>: 0.050 % | GCA_000073005.1 |

List of organisms from which reads were removed prior to assembly with Trinity. Quality-controlled reads were mapped to the genomes of listed organisms and reads that mapped to any of the references were removed.

We assembled separate transcriptomes for male and female *E. suzannae* whole adult wasps with the remaining unmapped reads using Trinity v2.6.6 with default settings [37]. Transcript abundance was then estimated for each with kallisto using the “align_and_estimate_abundance.pl” command bundled with Trinity [38]. Transcripts with an estimated abundance below 0.5 transcripts per million (TPM) were removed from both assemblies as these may be lowly expressed isoforms of other transcripts, poorly assembled or chimeric transcripts, or are simply contaminants and not from *Encarsia* [39, 40]. Next, TransDecoder v5.5.0 [41] was used to predict coding sequences within the remaining transcripts in each assembly and translate those coding sequences into predicted protein sequences with a minimum amino acid length of 67. Similar protein-coding sequences were then clustered using CD-HIT v4.6.8 [42, 43] with a threshold of 95% amino acid identity, and the longest protein isoform was assigned as the representative sequence for that cluster. The final assemblies are presented as the nucleotide sequence of the representative protein for each cluster. For a comprehensive list of the number of reads or transcripts at each step in the pipeline, see Table 2.

**Table 2:**
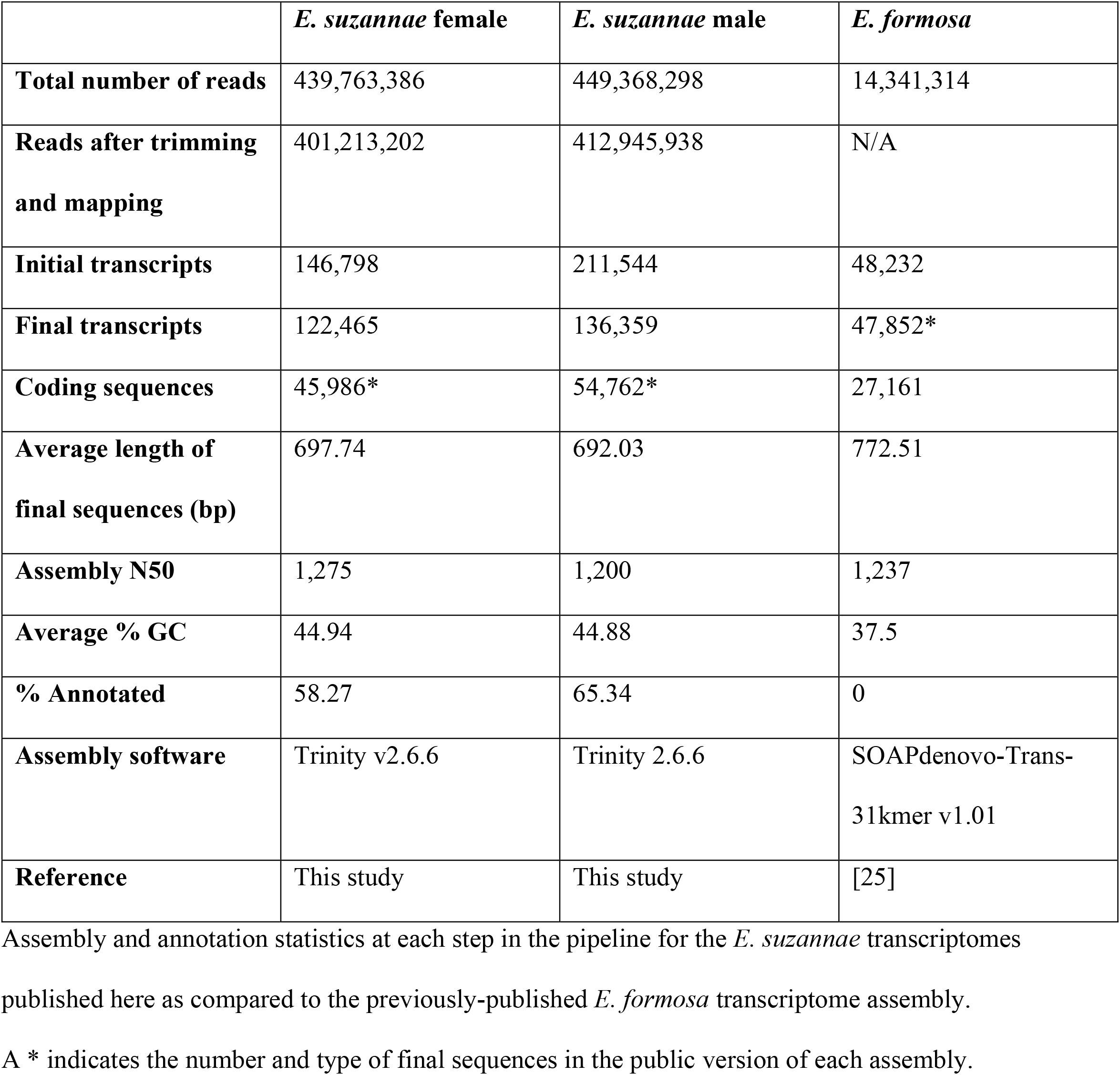
*E. suzannae* transcriptome read and transcript statistics.

### Quality control and data validation

Along with our mapping-and-removal approach to limit contamination while enriching for *Encarsia* reads prior to assembly, we also utilized additional methods to improve the quality of our assemblies. First, to comply with NCBI’s Transcriptome Shotgun Assembly (TSA) database requirements, we removed all coding sequences below 200bp. Furthermore, a blastn of remaining sequences against NCBI’s vector database was conducted to identify contaminating sequences and synthetic RNA spike-in controls, and hits with 100% nucleotide identity to vector sequences were removed from each assembly [44]. Prior to submission, any remaining coding sequences flagged by NCBI’s contamination check as sequencing vectors or contaminants were also removed. In total, 71 and 109 contaminating sequences were removed from the female and male assemblies, respectively.

The final assemblies were then assessed for completeness using BUSCO v5.3.2 in protein mode against the hymenoptera_odb10 reference lineage (v2020-08-05) [45, 46]. The female and male assemblies were found to possess, respectively, 82.1% and 82.6% of 5,991 complete orthologs identified as single-copy and nearly universal within the order Hymenoptera (present in >90% of species tested). This indicates a high level of completeness for both *E. suzannae* transcriptomes, although with varying degrees of duplication (shown in Table 3).

**Table 3:** Prediction of *E. suzannae* transcriptome assembly completeness using BUSCO.

| BUSCO results | Male *E. suzannae* |  | Female *E. suzannae* |  |
| --- | --- | --- | --- | --- |
|  | BUSCOs<br>present | Percent of<br>total | BUSCOs<br>present | Percent of<br>total |
| Complete BUSCOs | 4953 | 82.6 % | 4915 | 82.1 % |
| Complete single-copy BUSCOs | 3591 | 59.9 % | 4492 | 75.0 % |
| Complete duplicated BUSCOs | 1362 | 22.7 % | 423 | 7.1 % |
| Fragmented BUSCOs | 279 | 4.7 % | 280 | 4.7 % |
| Missing BUSCOs | 759 | 12.7 % | 796 | 13.2 % |
| Total BUSCO groups searched | 5991 | 100 % | 5991 | 100 % |

Assessment of assembly completeness using BUSCO v5.3.2 to search assembled proteins against a database of proteins identified as Hymenopteran BUSCOs. All BUSCO groups searched were determined to be present in single copy in >90% of Hymenopteran species tested; therefore, a high number of complete single-copy BUSCOs indicates a comprehensive and non-redundant assembly [45].

One issue which we are unable to rectify with the currently available sequencing data is the presence of *E*. sp. nr. *emiratus* transcripts within the male *E. suzannae* assembly. As mentioned above, haploid male *E. suzannae* eggs are laid into *Eretmocerus* pupae, and since this host does not have a sequenced genome (in contrast to *B. tabaci*), we could not apply the same mapping-and-removal approach to *E*. sp. nr. *emiratus*. This may be at least partly responsible for the elevated number of total sequences and duplicated BUSCOs in the *E. suzannae* male assembly compared to the female assembly (see Tables 2 and 3), but there are likely other contributing factors. Due to the relatedness of *Encarsia* and *Eretmocerus*, we are unable to differentiate sequences originating from those organisms at the read or assembled transcript level without a reference genome for either. However, we are confident that the abundance of *Eretmocerus* transcripts in the male assembly is low and many may have been removed from the assembly during the transcript abundance filtering step. This is evidenced by the very low *Eretmocerus* biomass present in/on fully emerged adult *E. suzannae* (larval *Encarsia suzannae* void their gut prior to pupation [47]), and using the average abundance of *B. tabaci* reads in either assembly as a proxy for *E*. sp. nr. *emiratus* reads suggests an abundance of around 5% for *Eretmocerus* (Table 1).

### Annotation

The male and female *E. suzannae* assemblies are available as unannotated coding sequences at NCBI’s TSA database under the accession numbers GJLB00000000 and GJLI00000000, respectively. Here, we also provide annotation information for both assemblies from multiple sources.

The final clustered proteins were annotated through the eggNOG-mapper v2 web-based pipeline using default settings to assign taxonomy to sequences and generate an annotation report with Gene Ontology (GO) terms, Pfam domains, KEGG pathway info, and other relevant information [48, 49]. Final proteins were also subjected to a search using DIAMOND with the “--very-sensitive” option [50] against NCBI’s non-redundant (nr) protein database (release 242.0) and a blastp search [51, 52] against a targeted database of well-annotated insect predicted proteomes consisting of *Nasonia vitripennis* Nvit_psr_1_1 (Genbank accession: GCA_009193385.2), *Trichogramma pretiosum* Tpre_2_0 (Genbank accession: GCA_000599845.3), and *Bemisia tabaci* MEAM1 (Genbank accession: GCA_001854935.1) using an e-value cutoff of 10^−5^. Although not closely related to *Encarsia, Bemisia* was included in the targeted insect database as its thorough annotation and presence as an outgroup may be useful in annotating proteins retained in *Encarsia* that *Nasonia* or *Trichogramma* may have lost. This database was also found to generate fewer hits labeled as “hypothetical” or “uncharacterized” when compared to a search against the nr protein database. The annotation results from each reference for both assemblies were pooled into a single Microsoft Excel spreadsheet (Additional File 1) and we have also provided an additional .fasta file for each assembly containing the final nucleotide sequences with sequence headers containing annotations from the blastp against the targeted insect database for ease of use (female: Additional file 2; male: Additional file 3).

Approximately 58% and 65% of female and male assembled proteins were annotated by one of the listed methods, with the characterization against NCBI’s nr database annotating the greatest number of proteins (26,155 female and 35,073 male), followed closely by the targeted insect database (24,478 female and 33,353 male). Some transcripts of note that were annotated in both the male and female assemblies are putative homologs to an array of insect sex-determination genes characterized in *Drosophila*. Homologs included *sex lethal* (*sxl*), the master regulator of the *Drosophila* sex-determination cascade, and some genes it regulates, including *transformer* (*tra), doublesex* (*dsx*), and *fruitless* (*fru*). *Sex lethal* controls the splicing of *tra* which itself is involved in the sex-specific splicing of *dsx* and *fru* [53]. In *Drosophila*, splicing by *tra* results in either male isoforms of *dsx* and *fru* or a female isoform of *dsx* and a truncated and untranslated female *fru* isoform. The different *dsx* isoforms are crucial for male and female somatic sexual development while *fru* appears to be key in the generation of male courtship behavior in *Drosophila* [54, 55]. We also searched the assemblies for homologs of *wasp overruler of masculinization* (*wom*) [56], which was identified in *N. vitripennis* as the instructor of sex determination via the activation of *tra* expression and autoregulation which, in turn, results in female development, but found none. However, we cannot rule out the presence of *wom* in *E. suzannae* as this gene in *N. vitripennis* is mainly transcribed in diploid (female) embryos prior to 7 hours post oviposition and is not expressed in adults, which we sampled for our transcriptome assemblies. We also did not find homologs of *complementary sex determiner* (*csd*), the instructor of sex determination in *Apis mellifera*.

Sex determination in the Chalcidoidea has been a matter of some speculation [57], but the presence of these transcripts provides insight into the nature of sex determination and development in *E. suzannae* and lays the foundation for understanding how the mechanisms of sexual development in *Encarsia* may interface with reproductive manipulation by *Cardinium*. Particularly applicable are cases of symbiont-induced parthenogenesis, in which unfertilized eggs are diploidized by the endosymbiont and biological females are produced [13, 58].

Furthermore, the identification of many transcripts harboring coding sequences annotated as putative venom proteins in both male and female *E. suzannae* transcriptomes is notable as these are believed to be important mechanisms used by female parasitoid wasps to enhance the survivability of their offspring. Venom proteins are diverse and are predicted to have a variety of impacts on the host undergoing parasitism, including immune system suppression, developmental arrest, lipid accumulation, apoptosis, and more [59]. In the case of *E. suzannae*, parasitism causes the whitefly host to undergo developmental arrest during a late nymphal stage. As arrest occurs regardless of wasp larva survival, it is possible that it is induced by venom injected into the whitefly during oviposition [15]. The presence of predicted proteins annotated as venom proteins in the male *E. suzannae* assembly is intriguing since only female wasps host feed and lay eggs into their host while adult males would seemingly have no need to express venom genes. It is unclear whether these putative proteins are actually venom genes expressed in male *E. suzannae* or if they were annotated as such due to the presence of domains similar to those found in venom proteins. Regardless, detecting putative venom proteins in *E. suzannae* provides more insight into how these wasps effectively parasitize their hosts; however, it should be noted that reliable identification of venom proteins will require additional experimental verification.

### Transcriptome comparisons

As stated above, the only other currently publicly available transcriptome of an *Encarsia* species belongs to *E. formosa* [26]; thus, limited comparisons can be made within this genus. An overview of all currently known *Encarsia* transcriptomes is shown in Table 2. Compared to the *E. formosa* transcriptome assembly, the male and female *E. suzannae* assemblies were generated from more initial reads and produced more transcripts pre-filtering, meaning they could be subject to more stringent transcript filtering than the *E. formosa* assembly. While the *E. formosa* assembly underwent limited post-assembly contaminant filtering, the *E. suzannae* assemblies utilized additional measures to 1) limit potential nonsense, low-abundance, and redundant transcripts through post-assembly filtering and processing, and 2) eliminate as many contaminants as possible prior to assembly via mapping-and-removal. Furthermore, the publicly available *E. formosa* assembly consists of full-length mRNA transcripts instead of coding sequences as seen in the *E. suzannae* assemblies [25]. After running TransDecoder on the *E. formosa* transcripts, only 27,161 coding sequences were predicted using a minimum length of 50 amino acids. This indicates that the female (45,986) and male (54,762) *E. suzannae* assemblies contain twice or nearly twice as many coding sequences compared to the *E. formosa* assembly, even though the *E. formosa* coding sequences were predicted with a shorter minimum protein size than *E. suzannae*.

Finally, OrthoVenn2 was used to determine orthologous groups between the predicted proteins in both *E. suzannae* assemblies presented in this paper and the *E. formosa* assembly published elsewhere [26, 60]. Using default settings and an e-value cutoff of 1e^-5^, 8,816 orthologs were found to be shared across all three transcriptomes, and a total of 22,015 orthologous groups were shared between male and female *E. suzannae* out of a total of 23,265 and 23,346 clusters, respectively (see Figure 1), indicating a high degree of similarity between the different sex assemblies but also showing the presence of over one thousand sex-specific protein clusters. It is also striking that female and male *E. suzannae* transcriptomes are equally similar to the *E. formosa* transcriptome despite the fact that *E. formosa* exists as an asexual species consisting of nearly all females (due to the presence of parthenogenesis-inducing *Wolbachia*) and its transcriptome therefore only reflects female individuals [61].

**Figure 1:**
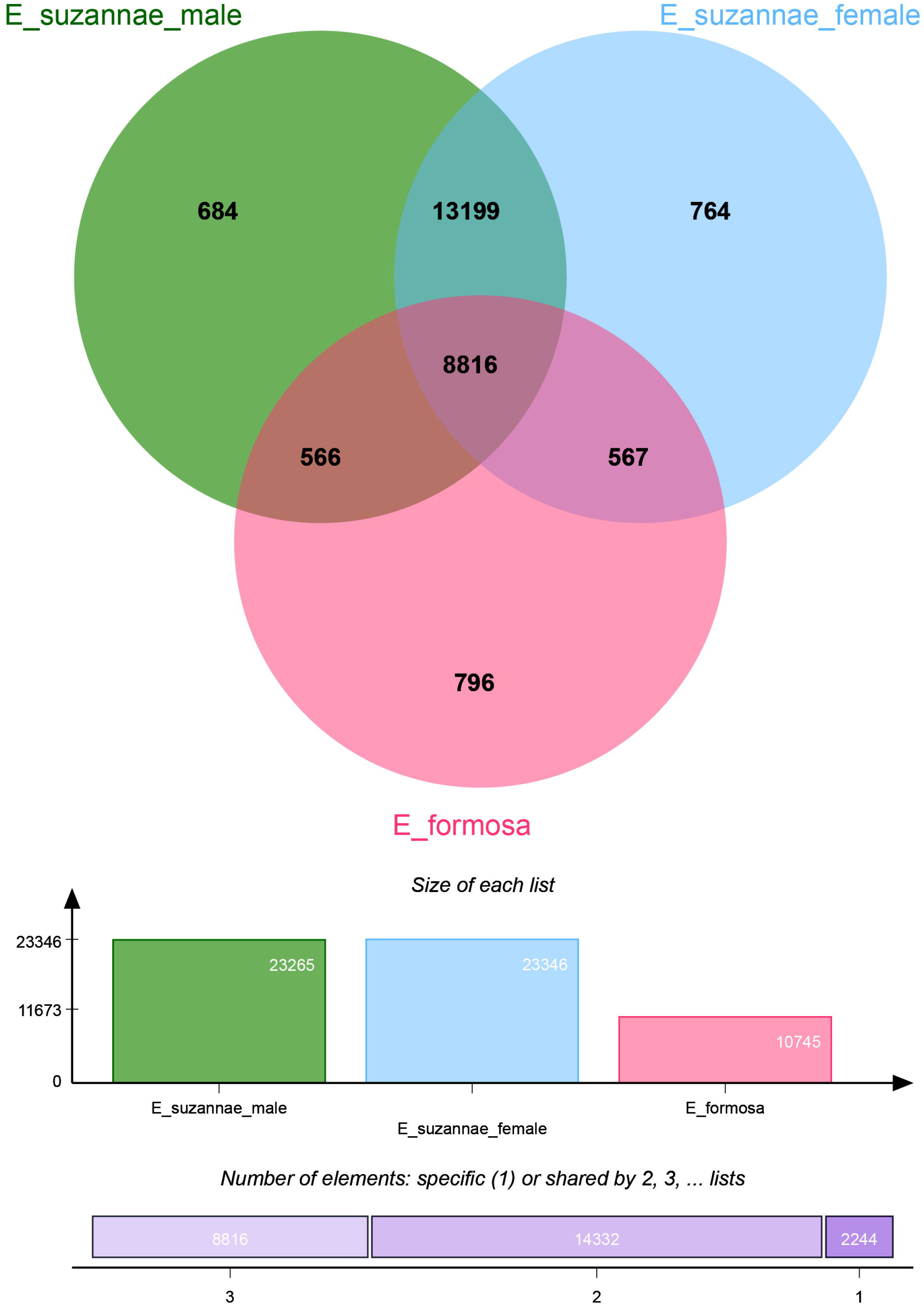
Orthologous groups between *E. formosa* females and male and female *E. suzannae* transcriptomes. The above figure shows an OrthoVenn2 diagram of orthologous groups between *E. formosa* females and male and female *E. suzannae* (e-value = 1e^-5^) [60]. TransDecoder using a minimum amino acid length of 50 was run on the *E. formosa* assemblies to obtain coding sequences and the resulting peptide sequence output (27,161 sequences) was tested against the predicted proteins from the male and female *E. suzannae* transcriptomes. The topmost Venn diagram depicts the number of shared orthologous protein clusters between the three transcriptomes. The middle bar graph depicts the total number of orthologous clusters present for each transcriptome, and the bottom graph shows (left to right) the number of clusters that were shared by all three transcriptomes, by any two transcriptomes, or were unique to only one of the three assemblies.

### Conclusion and re-use potential

We are confident that our assemblies are among the purest possible transcriptome representations of *E. suzannae* using the currently available data and assembly and filtering tools (for a list of all software and their versions utilized in this study, see Table 4). This study is also one of the first to present sex-specific transcriptome assemblies of a single insect species. In an organism such as *E. suzannae*, where males and females develop within different hosts, are impacted differently by endosymbiotic bacteria, and exhibit distinct behaviors, it is highly valuable to have available a reference database for both sexes to ensure more accurate studies when wasps of only one sex are used. Furthermore, these assemblies greatly expand our host knowledge of the *Cardinium c*Eper1 CI system and pave the way for future studies exploring how this endosymbiont interacts with its *E. suzannae* host in causing CI. We also believe that these data will be a valuable tool to other researchers as a reference when studying the diverse members of the ecologically important genus *Encarsia* and other chalcidoid parasitic wasps, many of which have interesting biology and potential as pest biological control agents.

**Table 4:** Software and version specifications.

| Software | Usage | Version | Reference(s) |
| --- | --- | --- | --- |
| BBTools | BBDuk for read trimming; BBMap for read mapping | 37.36 | [32] |
| FastQC | Visualization of sequence quality | 0.11.9 | [33] |
| SAMtools | .bam file manipulation | 1.10 | [62] |
| Trinity | *De novo* transcriptome assembly | 2.6.6 | [37] |
| kallisto | Transcript abundance estimation | 0.46.2 | [38] |
| TransDecoder | Prediction of coding sequences | 5.5.0 | [41] |
| CD-HIT | Clustering similar protein sequences | 4.6.8 | [42, 43] |
| BUSCO | Assessing assembly completeness | 5.3.2 | [45] |
| eggNOG-mapper v2 | Annotation of assembled proteins | 2.1.6 | [48, 49] |
| Blast+ | Annotation of assembled proteins | 2.11.0 | [51] |
| Diamond | Annotation of assembled proteins | 2.0.4 | [50] |
| OrthoVenn2 | Orthologous protein group clustering and visualization | N/A | [60] |

## Availability of supporting data

*E. suzannae* female and male raw read data and unannotated assemblies were submitted to NCBI’s Sequence Read Archive (SRA) and Transcriptome Shotgun Assembly (TSA) databases under the BioProjects PRJNA737477 for male *E. suzannae* and PRJNA737478 for female *E. suzannae*. Detailed annotation information from multiple sources is provided in Additional file 1. Annotated female and male assemblies are available in FASTA format in Additional files 2 and 3, respectively. All raw sequencing data and the final assemblies from this study are publicly available.

## Additional files

**Additional file 1:** Annotation results for female and male *E. suzannae* from different annotation methods

**Additional file 2:** FASTA format file containing the female *E. suzannae* transcriptome as coding sequences with annotations in the sequence headers

**Additional file 3:** FASTA format file containing the male *E. suzannae* transcriptome as coding sequences with annotations in the sequence headers

### Abbreviations

BUSCO: Benchmarking Universal Single-Copy Orthologs
CI: Cytoplasmic incompatibility
GO: Gene Ontology
KEGG: Kyoto Encyclopedia of Genes and Genomes
NCBI: National Center for Biotechnology Information
SRA: Sequence Read Archive
TPM: Transcripts per million
TSA: Transcriptome Shotgun Assembly
VBCF: Vienna BioCenter Core Facilities

## Authors’ contributions

SSE and MSH conceived the experiments and provided supervision. DLS and SSE developed the analysis pipeline. ES, CS, and SEK performed experiments. DLS analyzed and visualized the data and wrote the draft manuscript. All authors wrote and edited the manuscript. SSE and MSH obtained funding.

## Acknowledgements

We would like to thank Manuel Kleiner for his constructive comments and advice regarding the filtering and assembly pipeline.

